# Pneumonia Detection with Semantic Similarity Scores

**DOI:** 10.1101/2021.10.14.464247

**Authors:** Rahil Gholamipoor, Nima Rafiee, Markus Kollmann

**Affiliations:** Department of Computer Science, Heinrich Heine University, Düsseldorf, Germany; Department of Biology, Heinrich Heine University, Düsseldorf, Germany

## Abstract

X-ray images have been widely used for medical diagnoses of cardiothoracic and pulmonary abnormalities due to its noninvasiveness. Advancement in computer-aided diagnostic technologies, such as deep supervised methods, can help radiologists with a reliable early treatment and reduce diagnosis time. Nevertheless, these methods are prone to the small number of labeled samples and are limited to a specific abnormality. In this paper we combined a self-supervised contrastive method with a Mahalanobis distance score to develope an abnormality detection method that uses only healthy images during the training procedure. We were able to outperform previous unsupervised methods for the task of Pneumonia detection. We show that representation learned by the self-supervised method improves the supervised tasks for Pneumonia detection.

## 1. INTRODUCTION

Chest X-ray has been used for medical screening in order for the detection of cardiothoracic and pulmonary abnormalities which are one of the causes of mortality worldwide. Radiologists widely use chest X-ray images to diagnose lung-related diseases such as pneumonia. A computer-aided diagnostic approach would be very helpful to allow radiologists to detect potential abnormalities in chest X-ray images for early care and treatment. Recently supervised deep learning approaches have achieved promising results in abnormality detection for these images. Hendrycks et al. [1] proposed the maximum value of posterior distribution from the classifier as a baseline method to detect anomalies and Liang et al. [2] improved performance using temperature scaling and input pre-processing. However these approaches [3] require large, annotated datasets for training which is not always feasible. Additionally, it is in general difficult to acquire enough supervised data for rare pathologies. To address these problems, many approaches have exploited unsupervised or semisupervised frameworks to use unlabeled data for extracting generalizable features in medical images [4, 5]. Among unsupervised approaches, reconstruction-based methods assume that anomalies cannot be represented and reconstructed accu-rately by a model trained only on normal data. However, in practice these models can also reconstruct abnormal samples fairly well and thus fail to detect them [5, 6]. To overcome this problem, Mao et al. [7] trained an autoencoder model to not only reconstruct the corresponding normal version of any input, but also estimate the uncertainty of reconstruction at each pixel to enhance the performance of anomaly detection. In [8], an autoencoder is trained while a constraint is additionally imposed on the lower dimensional representation of the data in which features of the same X-ray images under random data augmentations are invariant, while the features of different images are scattered.

Recently the effectiveness of self-supervised contrastive learning has been proven in different domains, e.g. the visual domain [9, 10], which enables learning of robust representations through unlabeled data. Azizi et al. [11] investigated the effect of self-supervised pre-training on the classification downstream task on the CheXpert dataset. Zhang et al. [12] improved on supervised-based pneumonia detection using a contrastive-based pre-training and leveraging image description as an extra modality. In this paper, we utilize a self-supervised contrastive method to construct an anomaly detection score based on Mahalanobis distance for anomaly detection. To the best of our knowledge, we achieved state-of-the-art results for anomaly detection among all methods that can be applied to unlabeled data.

## 2. METHOD

### 2.1. Contrastive Learning

Given unlabeled training data, self-supervised contrastive representation learning aims to train a feature extractor, *g_θ_*, to discriminate similar samples from dissimilar ones. Using image transformations that keep the semantics, each image is augmented twice, referred to as positives. The function *g_θ_* is optimized to pull semantically similar samples together while pushing away from other images, referred to as negatives. Assuming that (*x_i_*, *x_j_*) is a positive pair for the *i^th^* image from a batch of *N* images, *τ* is a scalar temperature parameter and 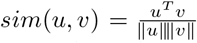 denotes the dot product between *l2* normalized *u* and *v* (i.e. cosine similarity). Contrastive learning minimizes the following loss for a positive pair of examples (*i, j*), referred to as Normalized Temperature-scaled Crossentropy (NT-Xent):

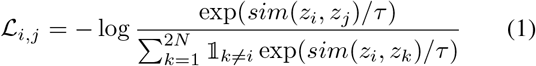

where 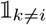 is an indicator function evaluating to 1 iff *k* ≠ *i*. *Z_i_* denotes the output feature of the contrastive layer. Intuitively, this loss is the log loss of a (2N)-way softmax-based classifier that tries to classify *x_j_* as a positive sample for *x_i_*. One can define the contrastive feature *z*(*x*) directly from the encoder *g_θ_*, i.e., *z*(*x*) = *g_θ_*(*x*) [10], or apply an additional projection layer *f_ϕ_*, i.e., *z*(*x*) = *f_ϕ_*(*g_θ_*(*x*)) [9]. The contrastive loss (Eq.1) can be minimized by different mechanisms that differ in how the negative samples are maintained. Chen et al. [9] takes negatives from the same batch but it requires a large batch size to provide a large set of negative pairs. Alternatively, Eq.1 can be minimized with sufficient number of negative pairs without using large batch sizes by maintaining negatives in a queue [10]. The encoded representations of the current mini-batch are enqueued while the oldest are dequeued. Unlike [9] in which only one encoder is used, following [10] we use two encoders, a query encoder and a slowly progressing key encoder, implemented as a momentum-based moving average of the query encoder.

### 2.2. Score Function for Anomaly Detection

#### Mahalanobis distance-based confidence score

We use Mahalanobis distance on feature space *h*(*x*) of the trained contrastive encoder as a score function for anomaly detection. Mahalanobis distance achieved promising results for supervised anomaly detection. Lee et al. [13] shows that with a well-trained softmax classifier, applying Mahalanobis distance on feature space using the class means and the feature covariance matrix can reach the state of the art results on supervised anomaly detection. To measure the Mahalanobis distance for a given test sample *x* first, we apply K-means clustering with *K* = 1 on the feature space *h*(*x*) of training data. This clustering helps to reduce computation time as we only compare the distance with the cluster mean. The anomaly score *s*(*x*) for a test sample *x* is given by the Mahalanobis distance

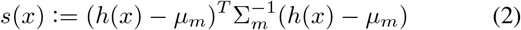

where *μ_m_* and Σ*_m_* are the mean and covariance of the feature vectors from the training data. The reason to use the Mahalanobis distance is to remove dominance of larger eigenvalues in euclidean distance metric as shown in [14] eigenvalues have an approximately inverse correlation with anomaly detection performance.

## 3. EXPERIMENTAL SETUP

### 3.1. Dataset

#### RSNA^1^

The Radiological Society of North America (RSNA) Pneumonia Detection Challenge dataset [15] is a publicly available dataset of frontal view chest radiographs. Each image was labeled as “Normal”, “No Opacity/Not Normal” or “Opacity”. The Opacity group consists of images with opacities suspicious for pneumonia, and images labeled “No Opacity/Not Normal” may have lung opacity, but no opacity suspicious for pneumonia. The RSNA dataset is a subset of the National Institutes of Health (NIH) Chest X-Ray dataset [16]. It contains 26, 684 X-rays with 8, 851 normal, 11, 821 no lung opacity/not normal and 6, 012 lung opacity.

### 3.2. Self-supervised Contrastive Training

Experiments were carried out using ResNet50 neural network architecture. Following [9], two fully connected layers are used to map the output of ResNet to a 128-dimensional embedding space where the contrastive loss is applied. We perform training on RSNA with initialization from ImageNet self-supervised pre-trained weights. We train at batch size 128 for 100 epochs using SGD optimiser. The temperature *τ* in Eq.(1) is set as 0.07. At training time, we apply the following augmentations: (1)a 224 × 224-pixel crop is taken from a randomly resized image (2) random rotation by an angle sampled from the uniform distribution *U*(−20, 20) (3) random horizontal flip with probability 0.5 (4) brightness and contrast adjustments.

### 3.3. Evaluation Methodology

We evaluate the results using Area Under the Receiver Operating Characteristic curve (AUROC), which has the advantage to be scale-invariant “it measures how well predictions are ranked, rather than their absolute values” and classificationthreshold-invariant “it measures how well anomaly samples are separated from the normal samples”.

## 4. EXPERIMENTAL RESULTS

### 4.1. Self-Supervised Anomaly Detection

For Mahalanobis distance, the highest performance achieved from the last layer, the output after the average pooling layer, before the MLP head [14]. On RSNA dataset, to detect anomalies we consider three different cases: “Normal” vs. “Opacity”; “Normal” vs. “No Opacity/Not Normal” and “Normal” vs. “all Opacity and No Opacity”. In Table 1, we compare our method with both supervised methods and unsupervised methods trained on only healthy images. We averaged AUROC values over 5 different train/test splits.

**Table 1.** OOD detection performance (%AUROC).

| Methods | Opacity | No Opacity | All |
| --- | --- | --- | --- |
| *Methods making use of label information* |  |  |  |
| Automated Abnormality Classification [17] | 0.980 | - | 0.949 |
| Pneumoni Detection using Radiomic Features[18] | 0.923 | - | - |
| ConVIRT [12] | - | - | 0.908 |
| *Unsupervised methods trained on normal samples* |  |  |  |
| UAE[7] | 0.89 | 0.78 | 0.83 |
| Deep Anomaly Detection[19] | 0.838 | 0.704 | 0.752 |
| Generative Adversarial one-class classifier[5] | 0.802 | - | 0.841 |
| Ours | **0.940** | **0.828** | **0.866** |

### 4.2. Pre-training and Label Efficiency of Multilabel Classification

In addition to self-supervised anomaly detection task, we evaluate the learned representaion by its performance in KNN accuracy and the impact it has on downstream task of multilable classification. For the pre-training task we use the same data split statistics as in [17] including 21152 training samples (14159 abnormal and 6993 normal samples). We use the same optimization config as for the anomaly detection task. The self-supervised pre-trained model achieved 1-NN accuracy of 79.01%. For the classification task, we replace the projection head of contrastive encoder with a classification head projecting the data into a one dimensional scalar value and fine-tune the whole model with binary cross entropy loss and same optimization config as in [17]. To see the effect of self-supervised pre-training, we start with small fraction of training data and compare model AUROC performance on test data for different case studies. Figure 3 shows that self-supervised pre-training can significantly help with label efficiency and causes a considerable performance improvement when we have a small fraction of labeled samples for the downstream task. We achieve AUROC score of 94.4%when fine-tuning with all labeled training data and AUROC score of 82.97%when using only 100 labeled samples which are selected randomly from training data.

**Fig. 1.**
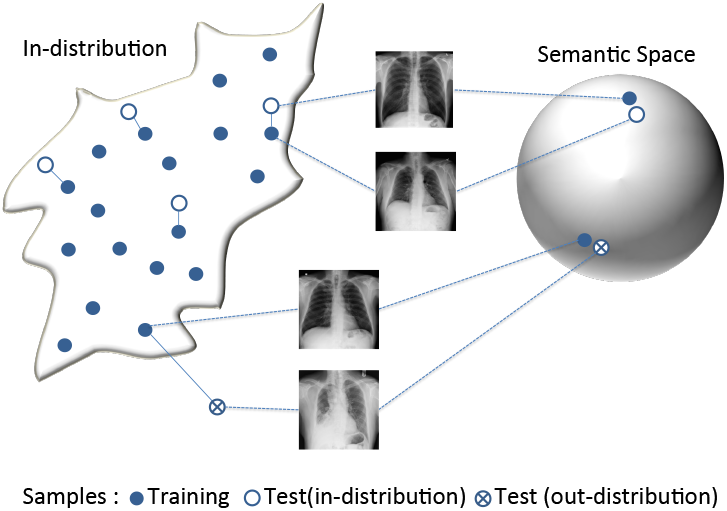
Illustration of mapping examples from the indistribution onto a unit-hypersphere. In this representation, feature vectors from the in-distribution are semantically similar if they approximately align and semantically diverse if they are separated by a large angle.

**Fig. 2.**
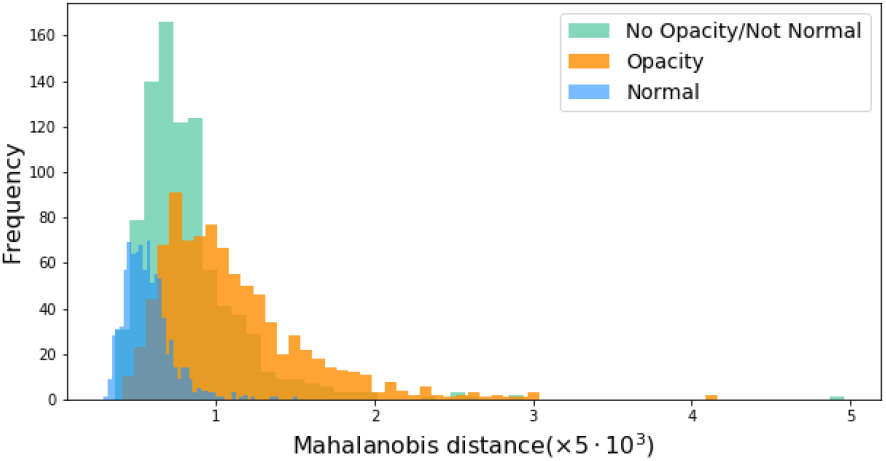
Distributions over the anomaly detection score trained only on Normal samples and applied to the test sets of Normal as in-distribution, Opacity and “No Opacity/Not Normal” as out-distributions.

**Fig. 3.**
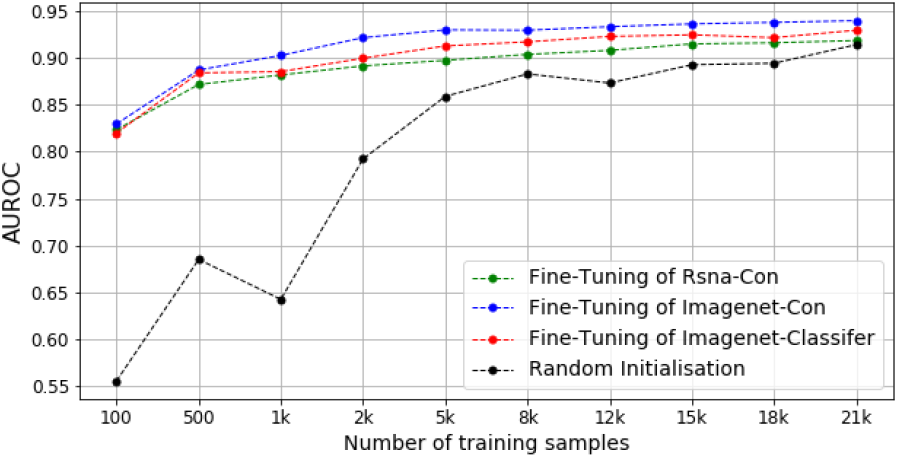
Self-supervised pre-training increases the downstream classification task performance with small fraction of training samples. Rsna-Con and Imagenet-Con are finetunings of models with different model initialisation in self-supervised pre-training as follows: randomly initialised and initialised with Imagenet. Imagenet-Classifier stands for fine-tuning an already trained imagenet classifier and Random Initialisastion is performing classification with random weight initialisation.

## 5. CONCLUSION

In this work, we proposed a self-supervised contrastive learning framework for X-ray anomaly detection trained only with the normal images to make our method future-ready for yet unknown anomalies. The self-supervised representations are highly effective for the task of anomaly detection in our framework. We define an anomaly detection score based on Mahalanobis distance applicable for detecting anomalies. We find that our approach outperforms all previous unsupervised methods on the RSNA pneumonia detection challenge dataset. This work may allow for improving radiology workflow and clinical decision-making.

## Footnotes

1 https://www.kaggle.com/c/rsna-pneumonia-detection-challenge/data

